## Supplementary Materials for "Removing reference bias and improving indel calling in ancient DNA data analysis by mapping to a sequence variation graph"

S21 Principal Component Analysis estimated with present-day West Eurasians from the Human Origins dataset with ancient samples aligned with either vg graph or bwa aln 31
S22 PCA of a subset of the Human Origins to examine the effect of read length in PC coordinates . . . 32

S23 Principal Component Analysis restricted to indels identified in Yamnaya processed with bwa and vg . . . 33

Table S1: Alternate allele fraction obtained after alignment of simulated data with vg, bwa aln and bwa mem, using different parameters and mapping quality filter thresholds.

| **aligner** | **mean alternate allele fraction and 95% CI** |
|---|---|
| vg graph q30 | 0.50001 [0.50000,0.50003] |
| vg graph q50 | 0.49988 [0.49984,0.49991] |
| vg graph q60 | 0.49933 [0.49924,0.49941] |
| vg linear q50 | 0.49555 [0.49510,0.49600] |
| bwa mem q50 | 0.48987 [0.48902,0.49073] |
| bwa aln -n 0.02 q25 | 0.49705 [0.49661,0.49750] |
| bwa aln -n 0.02 q30 | 0.48267 [0.48095,0.48438] |
| bwa aln -n 0.02 modreads q25 | 0.50015 [0.50014,0.50017] |
| bwa aln -n 0.02 modreads q30 | 0.50074 [0.50071,0.50077] |
| bwa aln -n 0.02 altref genome q25 | 0.50009 [0.50009,0.50009] |
| bwa aln -n 0.02 altref genome q30 | 0.49997 [0.49996,0.49999] |
| bwa aln -n 0.01 -o2 q25 | 0.49936 [0.49927,0.49945] |
| bwa aln -n 0.01 -o2 q30 | 0.49702 [0.49657,0.49747] |
| bwa aln n 0.01 -o2 modreads q25 | 0.50002 [0.50002,0.50003] |
| bwa aln n 0.01 -o2 modreads q30 | 0.50016 [0.50014,0.50017] |
| bwa aln n 0.01 -o2 altref genome q25 | 0.50011 [0.50011,0.50011] |
| bwa aln n 0.01 -o2 altref genome q30 | 0.50009 [0.50009,0.50009] |

Table S2: Comparison of the mean percentage of incorrectly mapped reads between vg alignment to the 1000GP graph, vg, bwa aln and bwa mem alignment to the linear reference at different mapping quality thresholds and alignment parameters. Error rates for the workflow which requires alignment to two reference genomes are also shown ('altref_genome').

| | | | | **mean error REF (%)** | | | |
|---|---|---|---|---|---|---|---|
| **mapping quality threshold** | **vg graph** | **vg linear** | **bwa mem** | **bwa aln -n0.02** | **bwa aln -n0.01 -o 2** | **bwa aln -n0.02 altref genome** | **bwa aln -n0.01 -o 2 altref genome** |
| **q>0** | 0.00714 | 0.00694 | 0.00315 | 0.00168 | 0.00168 | NA | NA |
| **q>=25** | 0.00018 | 0.00019 | 0.00024 | 0.00005 | 0.00002 | 0.00108 | 0.00062 |
| **q>=30** | 0.00015 | 0.00017 | 0.00016 | 0.00002 | 0.00001 | 0.00053 | 0.00054 |
| **q>=50** | 0.00012 | 0.00013 | 0.00007 | NA | NA | NA | NA |
| **q>=60** | 0.00011 | 0.00012 | 0.00004 | NA | NA | NA | NA |
| | | | | **mean error ALT (%)** | | | |
| | **vg graph** | **vg linear** | **bwa mem** | **bwa aln -n0.02** | **bwa aln -n0.01 -o 2** | **bwa aln -n0.02 altref genome** | **bwa aln -n0.01 -o 2 altref genome** |
| **q>0** | 0.01231 | 0.06035 | 0.02824 | 0.00980 | 0.00971 | NA | NA |
| **q>=25** | 0.00044 | 0.00191 | 0.00309 | 0.00054 | 0.00025 | 0.00051 | 0.00024 |
| **q>=30** | 0.00037 | 0.00162 | 0.00205 | 0.00024 | 0.00020 | 0.00021 | 0.00020 |
| **q>=50** | 0.00025 | 0.00110 | 0.00087 | NA | NA | NA | NA |
| **q>=60** | 0.00021 | 0.00097 | 0.00056 | NA | NA | NA | NA |
| | | | | **mean error All (%)** | | | |
| | **vg graph** | **vg linear** | **bwa mem** | **bwa aln -n0.02** | **bwa aln -n0.01 -o 2** | **bwa aln -n0.02 altref genome** | **bwa aln -n0.01 -o 2 altref genome** |
| **q>0** | 0.00972 | 0.03360 | 0.01563 | 0.00571 | 0.00569 | NA | NA |
| **q>=25** | 0.00031 | 0.00105 | 0.00165 | 0.00030 | 0.00013 | 0.0008 | 0.00043 |
| **q>=30** | 0.00026 | 0.00089 | 0.00109 | 0.00012 | 0.00011 | 0.00037 | 0.00037 |
| **q>=50** | 0.00019 | 0.00061 | 0.00046 | NA | NA | NA | NA |
| **q>=60** | 0.00016 | 0.00054 | 0.00029 | NA | NA | NA | NA |

Table S3: Comparison of the percentage of incorrectly aligned microbial reads of different lengths (30 to 100 bp) to the human reference genome between bwa aln, bwa mem and vg graph.

| **Read length** | **bwa aln -n0.01 -o2** | | **bwa aln -n0.02** | | **bwa mem** | | | **vg graph** | | |
|---|---|---|---|---|---|---|---|---|---|---|
| | **q>=25** | **q>=30** | **q>=25** | **q>=30** | **q>=30** | **q>=50** | **q>=60** | **q>=30** | **q>=50** | **q>=60** |
| **30** | 2.372 | 1.329 | 1.94 | 0.897 | 0.007 | 0.001 | 0.001 | 2.31 | 0.644 | 0.055 |
| **35** | 0.582 | 0.253 | 0.525 | 0.196 | 0.041 | 0.017 | 0.007 | 1.194 | 0.12 | 0.014 |
| **40** | 0.217 | 0.06 | 0.207 | 0.05 | 0.063 | 0.044 | 0.034 | 0.66 | 0.054 | 0.01 |
| **45** | 0.219 | 0.076 | 0.085 | 0.022 | 0.146 | 0.094 | 0.061 | 0.409 | 0.017 | 0.003 |
| **50** | 0.146 | 0.06 | 0.077 | 0.027 | 0.157 | 0.087 | 0.069 | 0.328 | 0.034 | 0.008 |
| **55** | 0.115 | 0.05 | 0.115 | 0.05 | 0.212 | 0.153 | 0.102 | 0.268 | 0.045 | 0.023 |
| **60** | 0.071 | 0.02 | 0.071 | 0.02 | 0.218 | 0.146 | 0.122 | 0.244 | 0.057 | 0.031 |
| **65** | 0.08 | 0.022 | 0.022 | 0.004 | 0.246 | 0.174 | 0.12 | 0.239 | 0.077 | 0.043 |
| **70** | 0.036 | 0.009 | 0.009 | 0 | 0.326 | 0.234 | 0.188 | 0.256 | 0.111 | 0.074 |
| **75** | 0.014 | 0.002 | 0.003 | 0 | 0.33 | 0.217 | 0.168 | 0.26 | 0.128 | 0.072 |
| **80** | 0.004 | 0.001 | 0.004 | 0.001 | 0.417 | 0.286 | 0.209 | 0.281 | 0.153 | 0.099 |
| **85** | 0 | 0 | 0 | 0 | 0.442 | 0.299 | 0.209 | 0.291 | 0.165 | 0.118 |
| **90** | 0.002 | 0 | 0 | 0 | 0.477 | 0.316 | 0.218 | 0.326 | 0.196 | 0.154 |
| **95** | 0.001 | 0.001 | 0.001 | 0.001 | 0.539 | 0.378 | 0.269 | 0.343 | 0.21 | 0.153 |
| **100** | 0 | 0 | 0 | 0 | 0.54 | 0.374 | 0.246 | 0.353 | 0.248 | 0.196 |

Table S4: Detailed description of the ancient DNA data processed in the present study.

| Set | Publication | SampleID | Type | Treatment | Reported coverage | Geographical Region |
|---|---|---|---|---|---|---|
| 1 | Damgaard et al., 2018 | Yamnaya | Shotgun | untreated | 13.6 | Kazakhstan |
| 1 | | Botai | Shotgun | untreated | 25.2 | Kazakhstan |
| 2 | Martiniano et al., 2016 | 1489 | Shotgun | untreated | 0.6 | United Kingdom |
| 2 | | 3DT16 | Shotgun | untreated | 0.7 | United Kingdom |
| 2 | | 3DT26 | Shotgun | untreated | 1.1 | United Kingdom |
| 2 | | 6DT18 | Shotgun | untreated | 1.1 | United Kingdom |
| 2 | | 6DT21 | Shotgun | untreated | 1.2 | United Kingdom |
| 2 | | 6DT22 | Shotgun | untreated | 1.1 | United Kingdom |
| 2 | | 6DT23 | Shotgun | untreated | 0.7 | United Kingdom |
| 2 | | 6DT3 | Shotgun | untreated | 1.7 | United Kingdom |
| 2 | | NO3423 | Shotgun | untreated | 1.1 | United Kingdom |
| 3 | Schiffels et al., 2016 | 12880A | Shotgun | USER | 1.3 | United Kingdom |
| 3 | | 12881A | Shotgun | USER | 4.4 | United Kingdom |
| 3 | | 12883A | Shotgun | USER | 3.8 | United Kingdom |
| 3 | | 12884A | Shotgun | USER | 11.8 | United Kingdom |
| 3 | | 12885A | Shotgun | USER | 0.9 | United Kingdom |
| 3 | | 15558A | Shotgun | UDGhalf | 3.8 | United Kingdom |
| 3 | | 15569A | Shotgun | UDGhalf | 2.7 | United Kingdom |
| 3 | | 15570A | Shotgun | UDGhalf | 8.2 | United Kingdom |
| 3 | | 15577A | Shotgun | UDGhalf | 6.3 | United Kingdom |
| 3 | | 15579A | Shotgun | UDGhalf | 1.4 | United Kingdom |
| 4 | Posth et al., 2018 | CP21half | Capture | UDGhalf | 2.5 | Brazil |
| 4 | | CP22half | Capture | UDGhalf | 2.0 | Brazil |
| 4 | | CP29half | Capture | UDGhalf | 5.3 | Peru |
| 4 | | CP8half | Capture | UDGhalf | 7.7 | Peru |
| 4 | | CUN008 | Capture | UDGhalf | 2.2 | Peru |
| 4 | | LAP003 | Capture | UDGhalf | 0.3 | Brazil |
| 4 | | LAP004 | Capture | UDGhalf | 1.2 | Brazil |
| 4 | | LAP005 | Capture | UDGhalf | 2.1 | Brazil |
| 4 | | LAP006 | Capture | UDGhalf | 0.9 | Brazil |
| 4 | | LAP007 | Capture | UDGhalf | 0.7 | Brazil |
| 4 | | LAR001 | Capture | UDGhalf | 0.6 | Brazil |
| 4 | | LAR002 | Capture | UDGhalf | 0.4 | Brazil |
| 4 | | MOS001 | Capture | UDGhalf | 0.4 | Brazil |

Table S5: Number of mapped reads, genomic coverage and endogenous content after aligning sequencing reads with vg to the 1000 Genomes variation graph.

| sampleID | number of reads | vg graph | | | | | | | | |
|---|---|---|---|---|---|---|---|---|---|---|
| | | aligned_mapq30_rmdups | endogenous_q30 | coverage_q30 | aligned_mapq50_rmdups | endogenous_q50 | coverage_q50 | aligned_mapq60_rmdups | endogenous_q60 | coverage_q60 |
| Botai | 1865598223 | 421934878 | 22.617 | 11.8103X | 395242184 | 21.186 | 11.3872X | 383123781 | 20.536 | 11.1798X |
| Yamnaya | 2663552613 | 804045151 | 30.187 | 19.3594X | 758072353 | 28.461 | 18.5996X | 734846662 | 27.589 | 18.211X |
| 1489 | 81132543 | 22320156 | 27.511 | 0.5485X | 21344354 | 26.308 | 0.5296X | 20859215 | 25.710 | 0.5199X |
| 3DT16 | 62769508 | 28831835 | 45.933 | 0.6661X | 27452525 | 43.735 | 0.6394X | 26767136 | 42.644 | 0.6259X |
| 3DT26 | 184353713 | 48564256 | 26.343 | 1.1371X | 46335474 | 25.134 | 1.0956X | 45249933 | 24.545 | 1.0745X |
| 6DT18 | 97642161 | 40381525 | 41.357 | 1.0543X | 38905880 | 39.845 | 1.0208X | 38182006 | 39.104 | 1.0041X |
| 6DT21 | 91458699 | 47424630 | 51.854 | 1.152X | 45480536 | 49.728 | 1.111X | 44513919 | 48.671 | 1.0904X |
| 6DT22 | 114753712 | 46199468 | 40.260 | 1.1107X | 44218645 | 38.534 | 1.0708X | 43242540 | 37.683 | 1.0508X |
| 6DT23 | 116564616 | 25504242 | 21.880 | 0.6459X | 24434218 | 20.962 | 0.6237X | 23927824 | 20.528 | 0.6126X |
| 6DT3 | 111539058 | 68660843 | 61.558 | 1.6627X | 65793129 | 58.987 | 1.6028X | 64354962 | 57.697 | 1.5725X |
| NO3423 | 89138342 | 43629793 | 48.946 | 1.0175X | 41665842 | 46.743 | 0.9792X | 40673417 | 45.630 | 0.9596X |
| 12880A | 829809746 | 56290616 | 6.784 | 0.9581X | 52677856 | 6.348 | 0.9108X | 51310381 | 6.183 | 0.8903X |
| 12881A | 794714619 | 172620548 | 21.721 | 2.7813X | 161674460 | 20.344 | 2.6227X | 157181011 | 19.778 | 2.553X |
| 12883A | 773030552 | 153957911 | 19.916 | 2.591X | 145022936 | 18.760 | 2.4591X | 141268741 | 18.275 | 2.3996X |
| 12884A | 1321644764 | 432117158 | 32.695 | 7.5554X | 410492176 | 31.059 | 7.2076X | 400887929 | 30.333 | 7.0476X |
| 12885A | 497373675 | 38872627 | 7.816 | 0.6481X | 36119828 | 7.262 | 0.6131X | 35100206 | 7.057 | 0.5982X |
| 15558A | 1054098249 | 152745125 | 14.491 | 2.8399X | 144221781 | 13.682 | 2.6959X | 140341229 | 13.314 | 2.6284X |
| 15569A | 237401000 | 93134209 | 39.231 | 1.6552X | 87821202 | 36.993 | 1.5664X | 85337285 | 35.946 | 1.5244X |
| 15570A | 823653962 | 299364605 | 36.346 | 5.2394X | 284333699 | 34.521 | 4.9962X | 277262836 | 33.663 | 4.8806X |
| 15577A | 683617585 | 232774127 | 34.050 | 4.3753X | 221020584 | 32.331 | 4.1814X | 215405385 | 31.510 | 4.0866X |
| 15579A | 327301910 | 55597145 | 16.987 | 0.7743X | 51828213 | 15.835 | 0.727X | 49962384 | 15.265 | 0.7029X |
| CP21half | 29829944 | 6125209 | 20.534 | 0.1219X | 5540251 | 18.573 | 0.112X | 5279948 | 17.700 | 0.1073X |
| CP22half | 24599607 | 5372420 | 21.839 | 0.0997X | 4806622 | 19.539 | 0.0904X | 4559164 | 18.533 | 0.086X |
| CP29half | 26634662 | 13403728 | 50.324 | 0.2736X | 12541204 | 47.086 | 0.2578X | 12141458 | 45.585 | 0.2502X |
| CP8half | 33014602 | 18898577 | 57.243 | 0.3924X | 17536032 | 53.116 | 0.3669X | 16915888 | 51.238 | 0.3548X |
| CUN008 | 24348976 | 6701266 | 27.522 | 0.1367X | 5971646 | 24.525 | 0.1231X | 5636410 | 23.148 | 0.1165X |
| LAP003 | 34760577 | 1112049 | 3.199 | 0.0174X | 883129 | 2.541 | 0.0148X | 805044 | 2.316 | 0.0137X |
| LAP004 | 79550494 | 4897788 | 6.157 | 0.0825X | 4084170 | 5.134 | 0.0711X | 3767481 | 4.736 | 0.0661X |
| LAP005 | 101557239 | 9350735 | 9.207 | 0.1696X | 7954168 | 7.832 | 0.1479X | 7374831 | 7.262 | 0.1379X |
| LAP006 | 62623053 | 4189609 | 6.690 | 0.0644X | 3596808 | 5.744 | 0.0567X | 3361499 | 5.368 | 0.0533X |
| LAP007 | 51237871 | 3322559 | 6.485 | 0.0512X | 2744928 | 5.357 | 0.0437X | 2527904 | 4.934 | 0.0406X |
| LAR001 | 12623337 | 1853770 | 14.685 | 0.0289X | 1630946 | 12.920 | 0.0256X | 1538141 | 12.185 | 0.0241X |
| LAR002 | 10812910 | 1249986 | 11.560 | 0.0197X | 1076414 | 9.955 | 0.0172X | 1008331 | 9.325 | 0.0161X |
| MOS001 | 17259402 | 1164601 | 6.748 | 0.0171X | 996820 | 5.776 | 0.0148X | 933975 | 5.411 | 0.0139X |

Table S6: Number of mapped reads, genomic coverage and endogenous content after aligning sequencing reads with bwa aln to the human reference genome.

| sampleID | number of reads | bwa -n 0.02 | | | | | | bwa -n 0.01 -o 2 | | | | | |
|---|---|---|---|---|---|---|---|---|---|---|---|---|---|
| | | aligned_mapq25_rmdups | endogenous_q25 | coverage_q25 | aligned_mapq30_rmdups | endogenous_q30 | coverage_q30 | aligned_mapq25_rmdups | endogenous_q25 | coverage_q25 | aligned_mapq30_rmdups | endogenous_q30 | coverage_q30 |
| Botai | 1865598223 | 416223409 | 22.310 | 11.3479X | 410152233 | 21.985 | 11.2361X | 415164292 | 22.254 | 11.274X | 410749213 | 22.017 | 11.1976X |
| Yamnaya | 2663552613 | 811584947 | 30.470 | 19.1319X | 801807300 | 30.103 | 18.952X | 809244367 | 30.382 | 19.0093X | 802441463 | 30.127 | 18.8915X |
| 1489 | 81132543 | 22535412 | 27.776 | 0.5518X | 22138624 | 27.287 | 0.5428X | 22514582 | 27.750 | 0.5505X | 22256397 | 27.432 | 0.545X |
| 3DT16 | 62769508 | 28595404 | 45.556 | 0.6614X | 28220816 | 44.959 | 0.6531X | 28553146 | 45.489 | 0.6596X | 28323320 | 45.123 | 0.6547X |
| 3DT26 | 184353713 | 47957571 | 26.014 | 1.1258X | 47211710 | 25.609 | 1.1091X | 47913845 | 25.990 | 1.1234X | 47443675 | 25.735 | 1.1133X |
| 6DT18 | 97642161 | 40243643 | 41.215 | 1.0492X | 39968525 | 40.934 | 1.0425X | 40102223 | 41.071 | 1.0446X | 39952714 | 40.917 | 1.0412X |
| 6DT21 | 91458699 | 47102981 | 51.502 | 1.143X | 46659138 | 51.017 | 1.1326X | 46986468 | 51.375 | 1.1389X | 46724635 | 51.088 | 1.1331X |
| 6DT22 | 114753712 | 45849456 | 39.955 | 1.1022X | 45384992 | 39.550 | 1.0918X | 45753752 | 39.871 | 1.0987X | 45474863 | 39.628 | 1.0927X |
| 6DT23 | 116564616 | 25166305 | 21.590 | 0.6388X | 24866360 | 21.333 | 0.6315X | 25101512 | 21.534 | 0.6365X | 24925776 | 21.384 | 0.6324X |
| 6DT3 | 111539058 | 68144820 | 61.095 | 1.6488X | 67425973 | 60.451 | 1.6322X | 67992471 | 60.958 | 1.6433X | 67564411 | 60.575 | 1.6339X |
| NO3423 | 89138342 | 44498077 | 49.920 | 1.0355X | 43903918 | 49.254 | 1.0225X | 44405123 | 49.816 | 1.0318X | 44037034 | 49.403 | 1.0241X |
| 12880A | 829809746 | 58594810 | 7.061 | 1.0066X | 58508458 | 7.051 | 1.0052X | 58588180 | 7.060 | 1.0063X | 58522112 | 7.052 | 1.0052X |
| 12881A | 794714619 | 185542021 | 23.347 | 2.9876X | 185119809 | 23.294 | 2.9811X | 185574034 | 23.351 | 2.9877X | 185217661 | 23.306 | 2.9824X |
| 12883A | 773030552 | 162441204 | 21.014 | 2.7352X | 162132983 | 20.974 | 2.7303X | 162436150 | 21.013 | 2.7346X | 162187543 | 20.981 | 2.7308X |
| 12884A | 1321644764 | 451142252 | 34.135 | 7.8728X | 450344000 | 34.075 | 7.8596X | 451081278 | 34.130 | 7.87X | 450442538 | 34.082 | 7.8598X |
| 12885A | 497373675 | 40245318 | 8.092 | 0.6794X | 40172448 | 8.077 | 0.6782X | 40246069 | 8.092 | 0.6792X | 40188524 | 8.080 | 0.6784X |
| 15558A | 1054098249 | 159809718 | 15.161 | 2.9646X | 159401847 | 15.122 | 2.9575X | 159759312 | 15.156 | 2.9627X | 159448052 | 15.126 | 2.9574X |
| 15569A | 237401000 | 97393061 | 41.025 | 1.7286X | 97202437 | 40.944 | 1.7253X | 97378920 | 41.019 | 1.728X | 97239275 | 40.960 | 1.7257X |
| 15570A | 823653962 | 310627818 | 37.713 | 5.4285X | 310093622 | 37.649 | 5.4196X | 310588471 | 37.709 | 5.4267X | 310179713 | 37.659 | 5.4201X |
| 15577A | 683617585 | 242272982 | 35.440 | 4.5412X | 241843836 | 35.377 | 4.5337X | 242195248 | 35.428 | 4.5384X | 241880760 | 35.382 | 4.5331X |
| 15579A | 327301910 | 57857227 | 17.677 | 0.8094X | 57716758 | 17.634 | 0.8074X | 57855021 | 17.676 | 0.8093X | 57749295 | 17.644 | 0.8079X |
| CP21half | 29829944 | 6409908 | 21.488 | 0.1286X | 6377413 | 21.379 | 0.1279X | 6413170 | 21.499 | 0.1286X | 6394889 | 21.438 | 0.1283X |
| CP22half | 24599607 | 5695703 | 23.154 | 0.1064X | 5661549 | 23.015 | 0.1057X | 5698772 | 23.166 | 0.1064X | 5678398 | 23.083 | 0.106X |
| CP29half | 26634662 | 13862670 | 52.047 | 0.2831X | 13799520 | 51.810 | 0.2818X | 13870225 | 52.076 | 0.2833X | 13833883 | 51.939 | 0.2826X |
| CP8half | 33014602 | 19322936 | 58.528 | 0.4029X | 19145655 | 57.991 | 0.3992X | 19356554 | 58.630 | 0.4036X | 19234195 | 58.260 | 0.4012X |
| CUN008 | 24348976 | 7232279 | 29.703 | 0.1466X | 7198507 | 29.564 | 0.1459X | 7231444 | 29.699 | 0.1466X | 7205168 | 29.591 | 0.146X |
| LAP003 | 34760577 | 1096188 | 3.154 | 0.0184X | 1086514 | 3.126 | 0.0183X | 1098299 | 3.160 | 0.0185X | 1090573 | 3.137 | 0.0184X |
| LAP004 | 79550494 | 5184747 | 6.518 | 0.0898X | 5141548 | 6.463 | 0.0891X | 5191313 | 6.526 | 0.0899X | 5160397 | 6.487 | 0.0894X |
| LAP005 | 101557239 | 10078860 | 9.924 | 0.1854X | 10006121 | 9.853 | 0.1841X | 10087457 | 9.933 | 0.1856X | 10037619 | 9.884 | 0.1847X |
| LAP006 | 62623053 | 4346951 | 6.941 | 0.0687X | 4308189 | 6.880 | 0.068X | 4351282 | 6.948 | 0.0687X | 4323436 | 6.904 | 0.0683X |
| LAP007 | 51237871 | 3425557 | 6.686 | 0.0547X | 3400460 | 6.637 | 0.0543X | 3429966 | 6.694 | 0.0548X | 3411388 | 6.658 | 0.0545X |
| LAR001 | 12623337 | 2014020 | 15.955 | 0.0316X | 2001981 | 15.859 | 0.0314X | 2014625 | 15.960 | 0.0316X | 2005651 | 15.888 | 0.0315X |
| LAR002 | 10812910 | 1383937 | 12.799 | 0.0221X | 1376256 | 12.728 | 0.022X | 1384288 | 12.802 | 0.0221X | 1378604 | 12.750 | 0.022X |
| MOS001 | 17259402 | 1263689 | 7.322 | 0.019X | 1255509 | 7.274 | 0.0188X | 1264502 | 7.326 | 0.019X | 1258051 | 7.289 | 0.0189X |

Table S7: Comparison of run times, memory and storage between bwa and vg for the indexing of the linear human reference genome and the 1000GP graph, respectively. We also compare read alignment and bam generation with vg, bwa aln and bwa mem, using an uncompressed FASTQ file containing 10 million simulated 50 bp reads with various levels of deamination.

| **Process** | **Command** | **CPU time** | **Max Memory (Gb)** | **Storage (Gb)** |
|---|---|---|---|---|
| Indexing | bwa index hsGR37.fa | 02:11:04 | 4.5 | 5.2 |
| | vg index 1000GP_AF0.001 | 26:40:14 | 121.1 | 38.1 |
| Read alignment and bam file generation | vg map -w 1024 -k15 –surject-to bam | 06:17:56 | 37.3 | - |
| | bwa mem + samtools view -Sb | 00:13:04 | 5.4 | - |
| | bwa aln -t1 -l1024 -q15 -n0.01 -o 2 + bwa samse + samtools view -Sb | 01:16:16 | 4.6 | - |
| | bwa aln -t1 -l1024 -q15 -n0.01 + bwa samse + samtools view -Sb | 01:22:43 | 3.9 | - |
| | bwa aln -t1 -l1024 -q15 -n0.02 + bwa samse + samtools view -Sb | 00:48:48 | 4.6 | - |

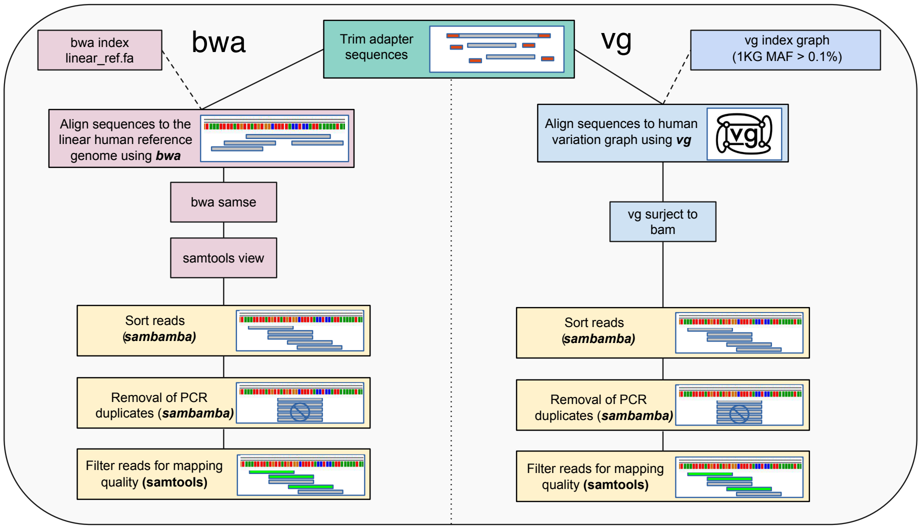

Figure S1: Diagram summarizing the data processing workflow.

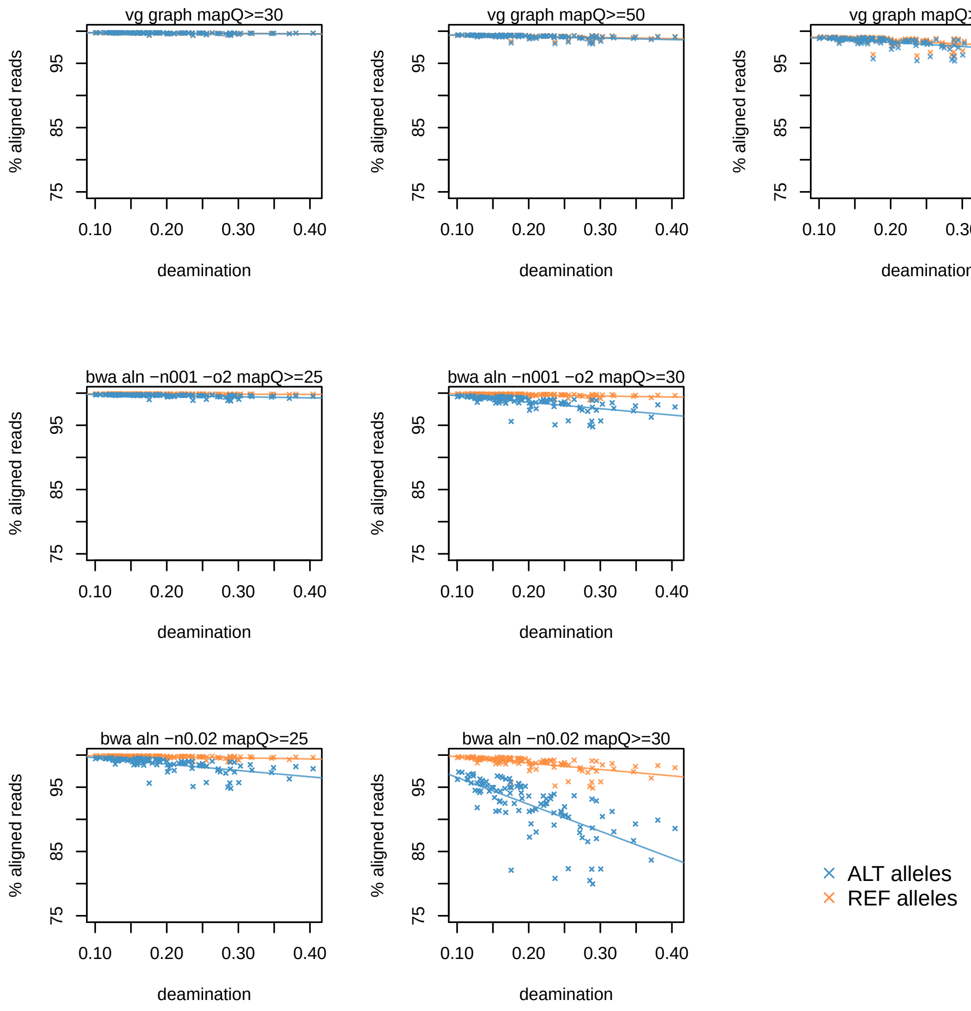

Figure S2: Comparison of the percentage of mapped reads in simulated data between vg graph and bwa aln (-n 0.02 and -n 0.01 -o 2) filtered with different mapping quality thresholds.

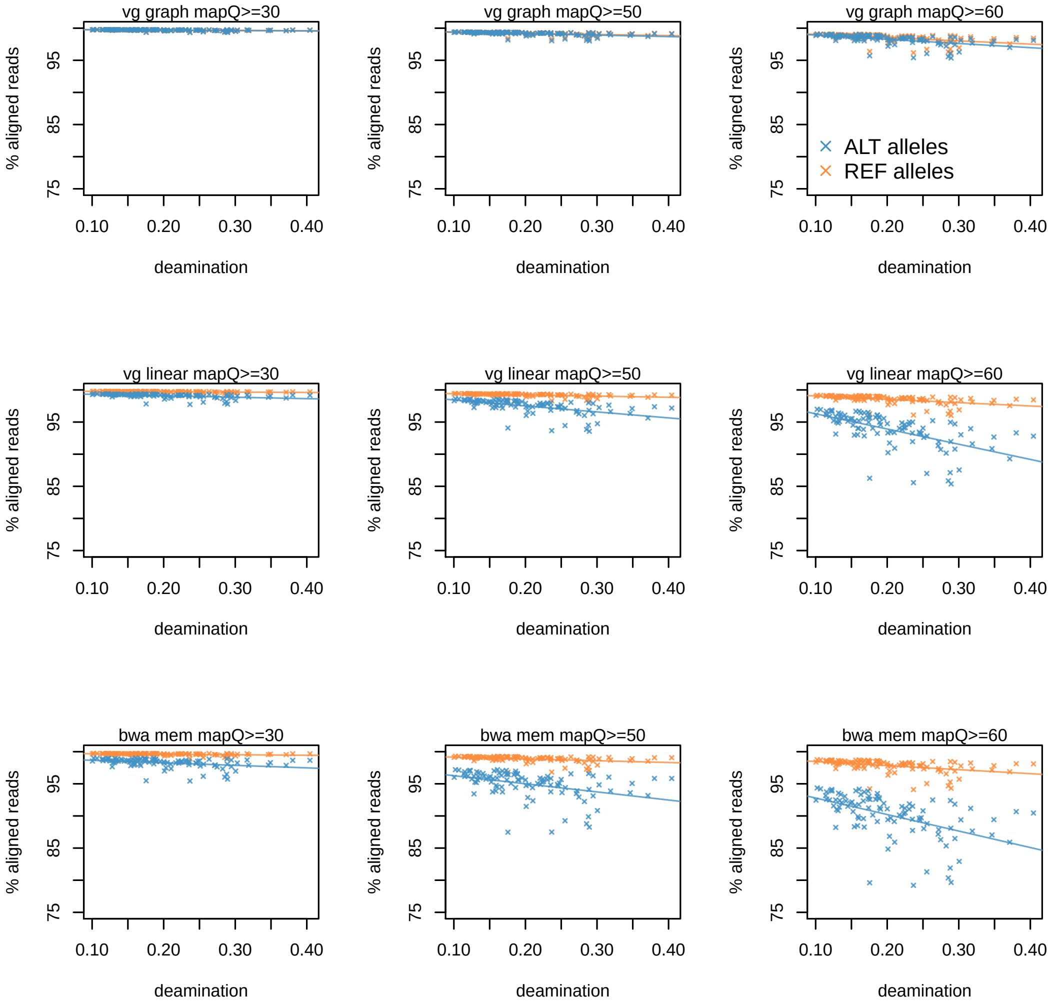

Figure S3: Comparison of the percentage of mapped reads in simulated data between vg graph, vg linear reference and bwa mem filtered with different mapping quality thresholds.

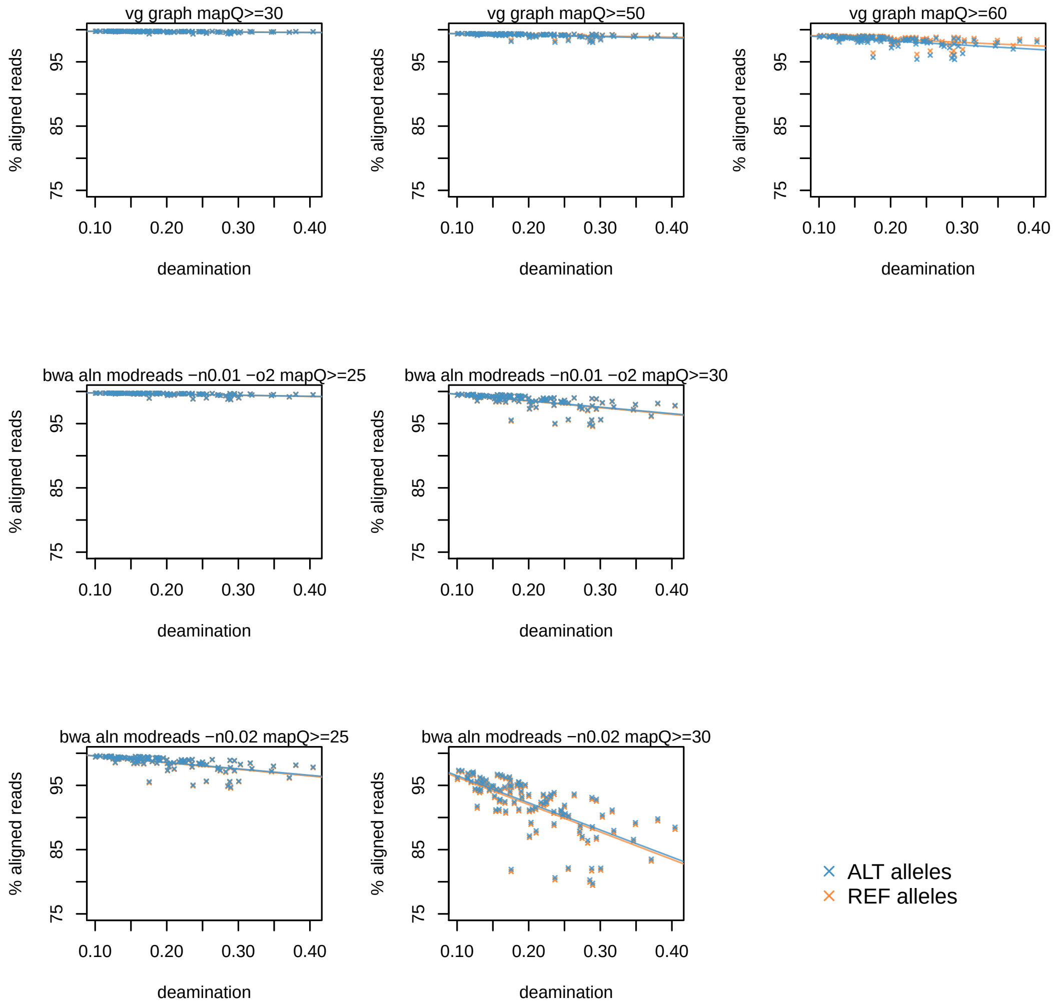

Figure S4: Comparison of the percentage of mapped reads in simulated data between vg graph and bwa aln (-n 0.02 and -n 0.01 -o 2) after processing with the Günther & Nettelblad method which modifies reads to remove reference bias.

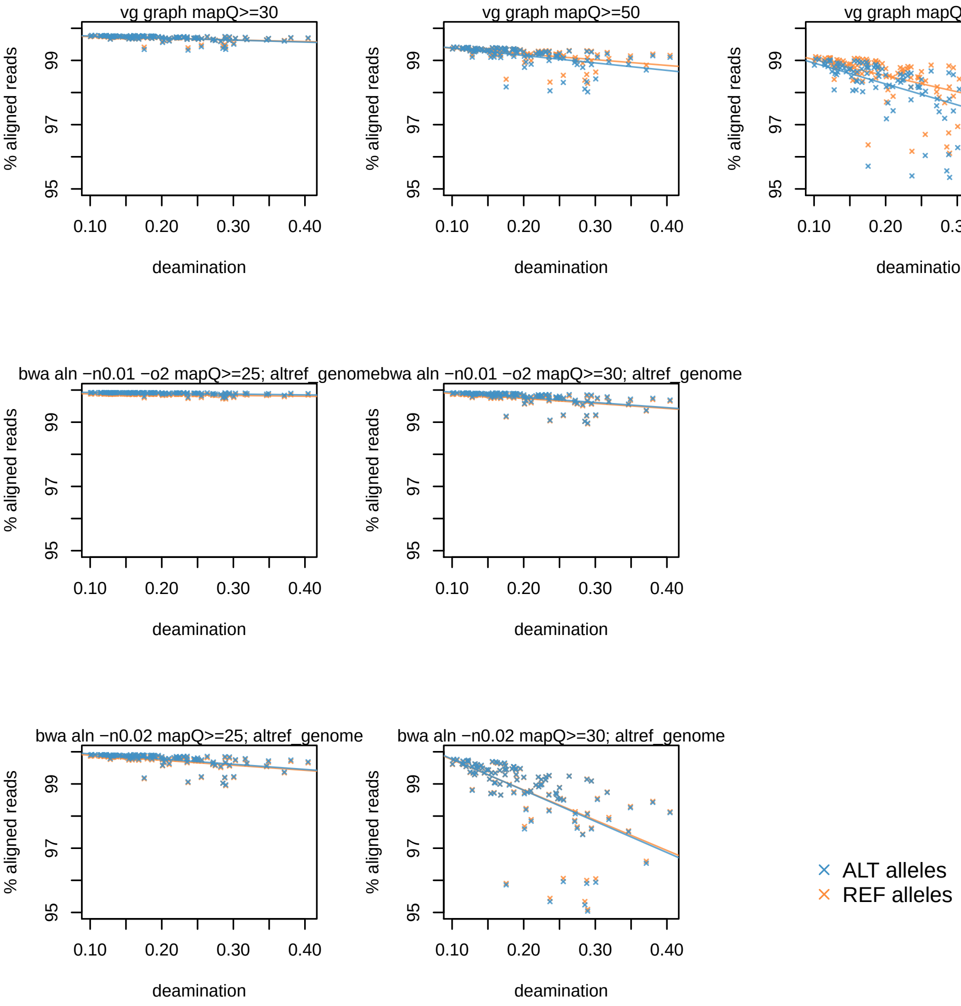

Figure S5: Comparison of the percentage of mapped reads in simulated data between vg and bwa aln (-n 0.02 and -n 0.01 -o 2) and post processing with the Peyrégne workflow, in which reads are mapped to two versions of the reference genome (one with the alternate and the other with the reference allele at Human Origins SNPs; 'altref_genome') in order to remove reference bias.

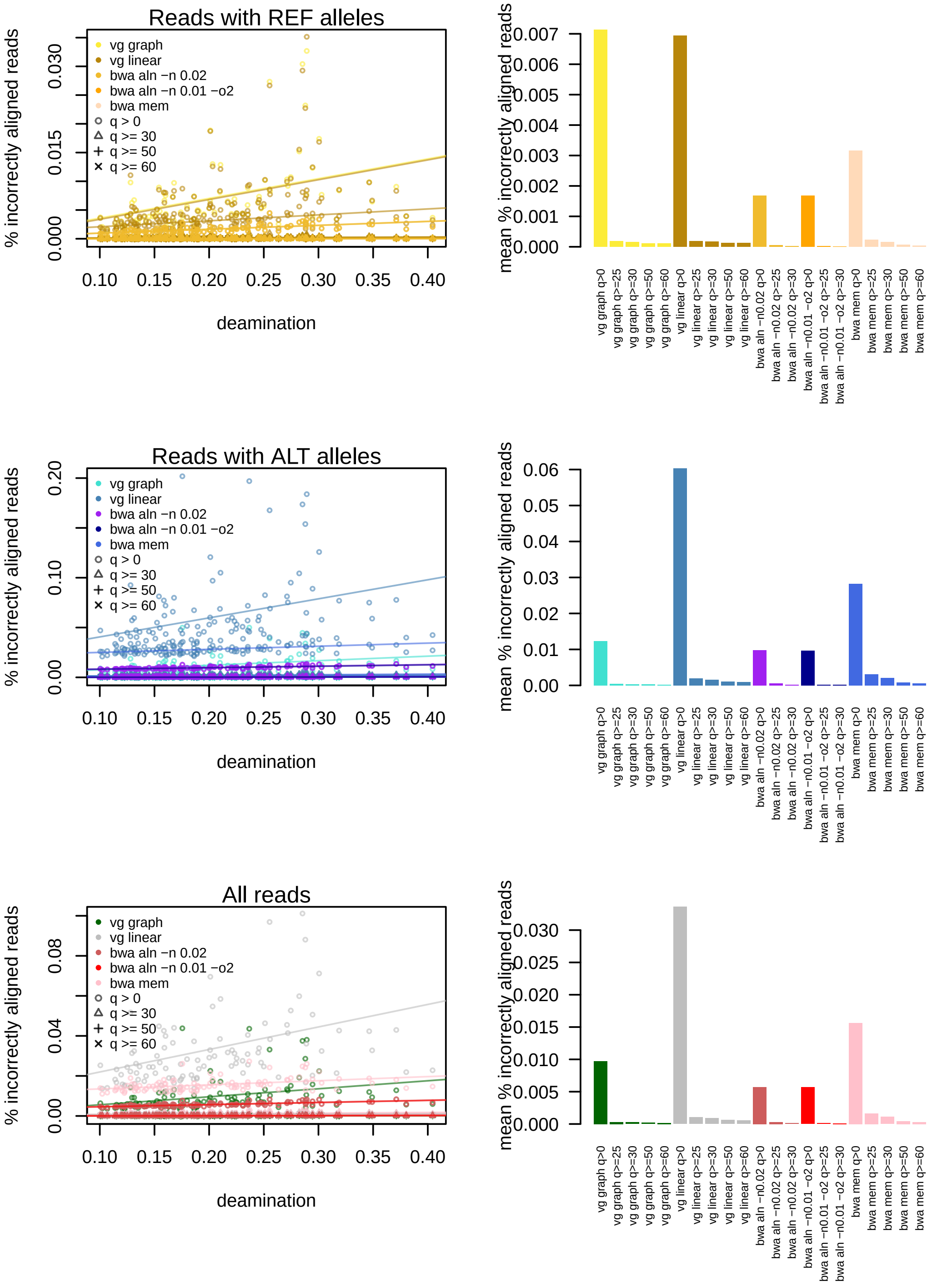

Figure S6: Comparison of the percentage of incorrectly mapped reads between bwa (aln and mem), vg and vg linear reference using different parameters and filtered with different mapping quality thresholds (q>0, q≥30, q≥50 and q≥60). The plots on the left show the percentage of incorrectly mapped reads at increasing deamination rates, and the barplots on the right indicate the mean for each aligner and imposed mapping quality threshold.

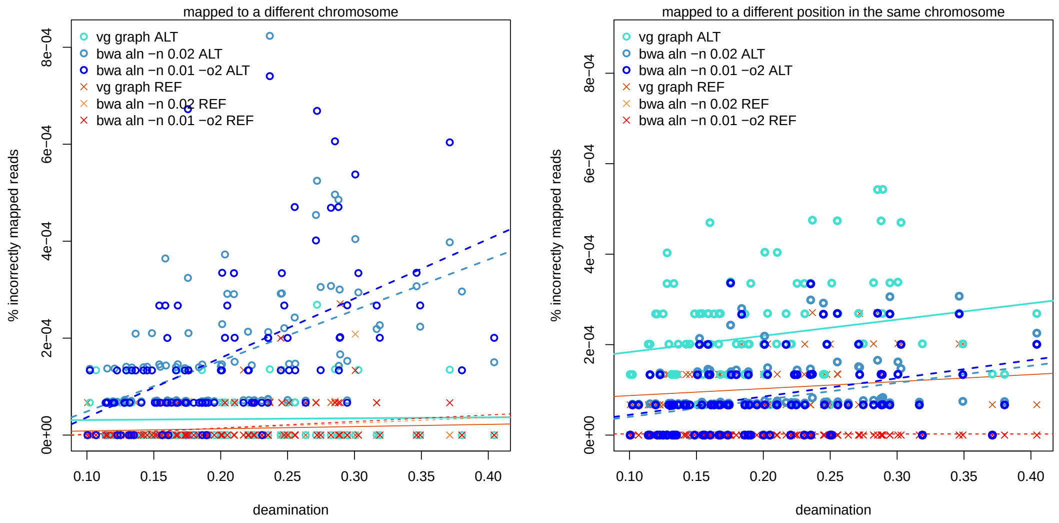

Figure S7: Relationship between deamination and alignment error in vg (q≥50) and bwa aln (q≥30) alignments. Reads were simulated from chr11. The left side plot shows the percentage of reads incorrectly mapped to a different chromosome, and the one on the right shows the percentage of reads incorrectly mapped to a different position of the same chromosome they were simulated from.

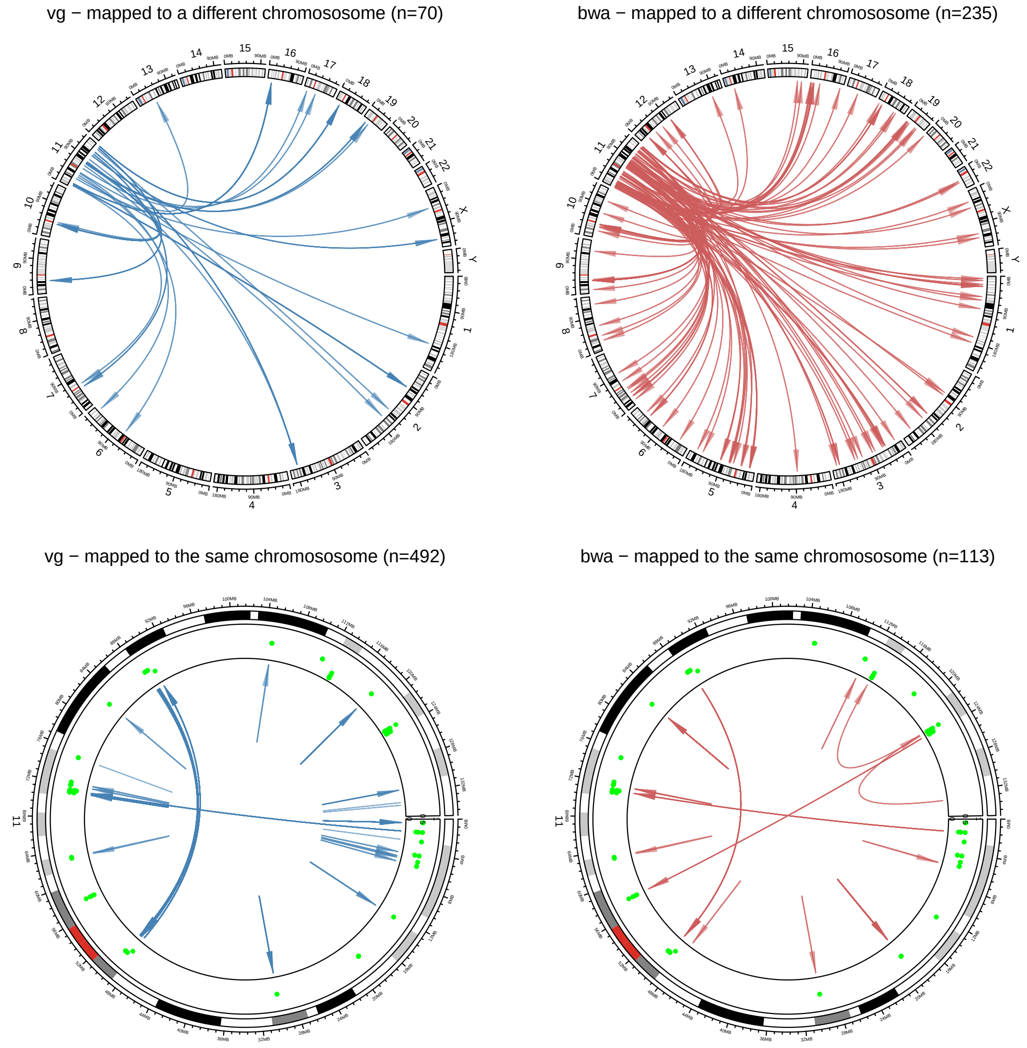

Figure S8: Circos plots comparing the types of errors observed in all the simulated reads aligned with bwa aln (-n 0.02 q$\geq$30) and vg graph (q$\geq$50). Alignment errors are represented by arrows, with each arrow connecting the region where reads were simulated from and the regions where they were aligned to. The top circos plots show cases where reads were mapped to different chromosomes, while the bottom plots show cases where a read was aligned to a different coordinate within chr11. The vast majority of mapping errors occur in reads overlapping regions with low mappability. Green points indicate overlaps with low mappability ($\leq$ 0.5) regions in the Mappability 50 track, downloaded from the UCSC table browser. A mappability of 0.5 means that a given read can be mapped equally well to two different places in the human reference genome. The numbers above each plot are the total error counts across all simulated individuals.

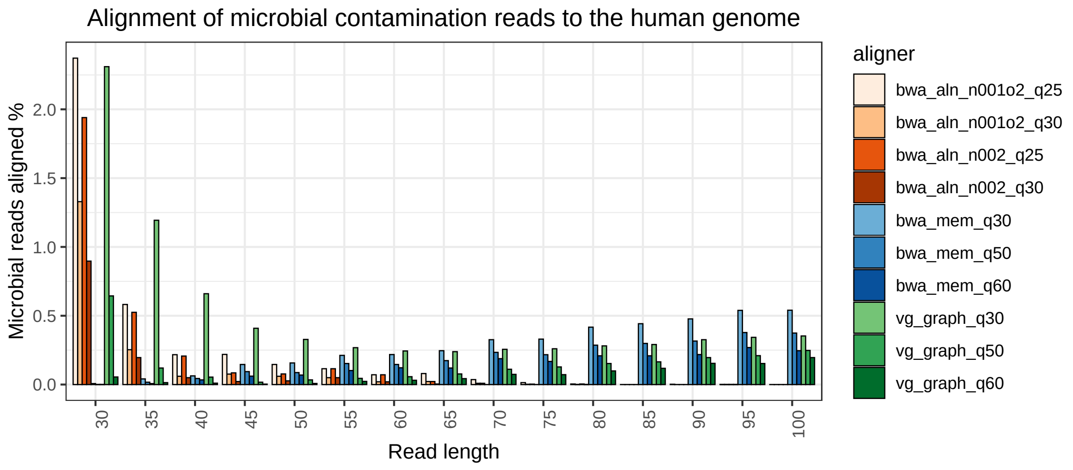

Figure S9: Comparison of spurious alignment of simulated microbial sequences of different read lengths (30-100bp) between bwa aln and bwa mem to the human reference genome and vg to the 1000GP graph.

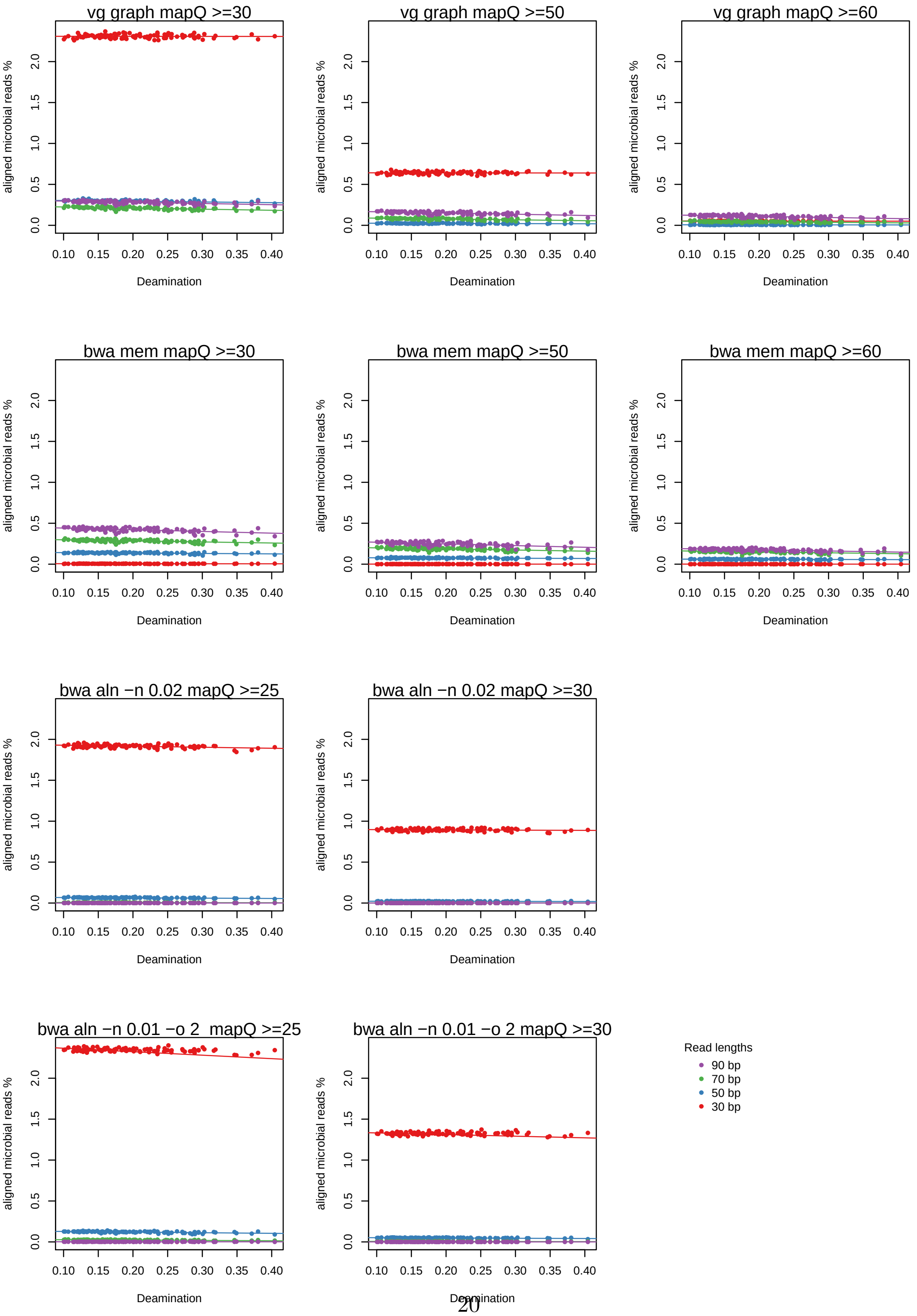

Figure S10: Investigation of the effect of deamination levels in the spurious alignment of simulated microbial sequences of different read lengths (30, 50, 70 and 90bp) in bwa aln, bwa mem and vg.

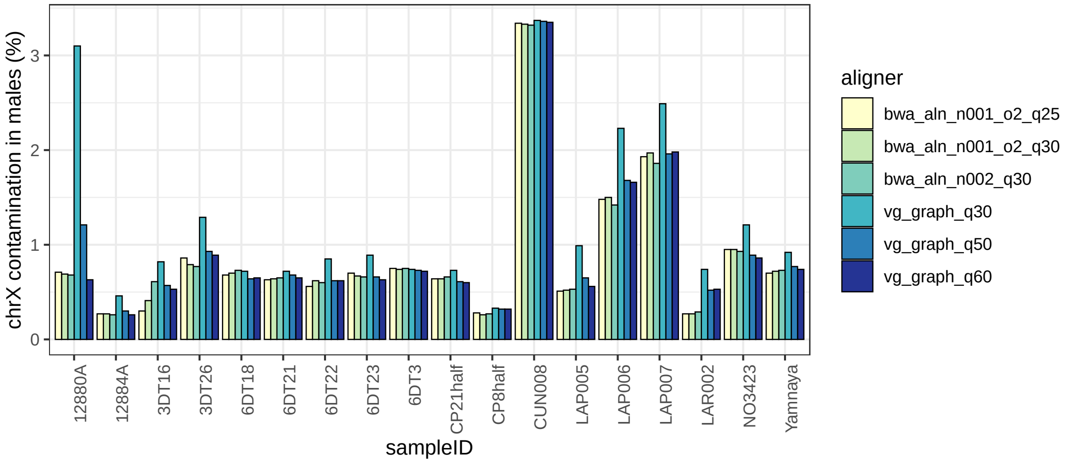

Figure S11: Comparison of the fraction of X-chromosome contamination estimates in male samples aligned with vg graph and bwa aln -0.02 and -n 0.01 -o2 and filtered using different mapping quality thresholds.

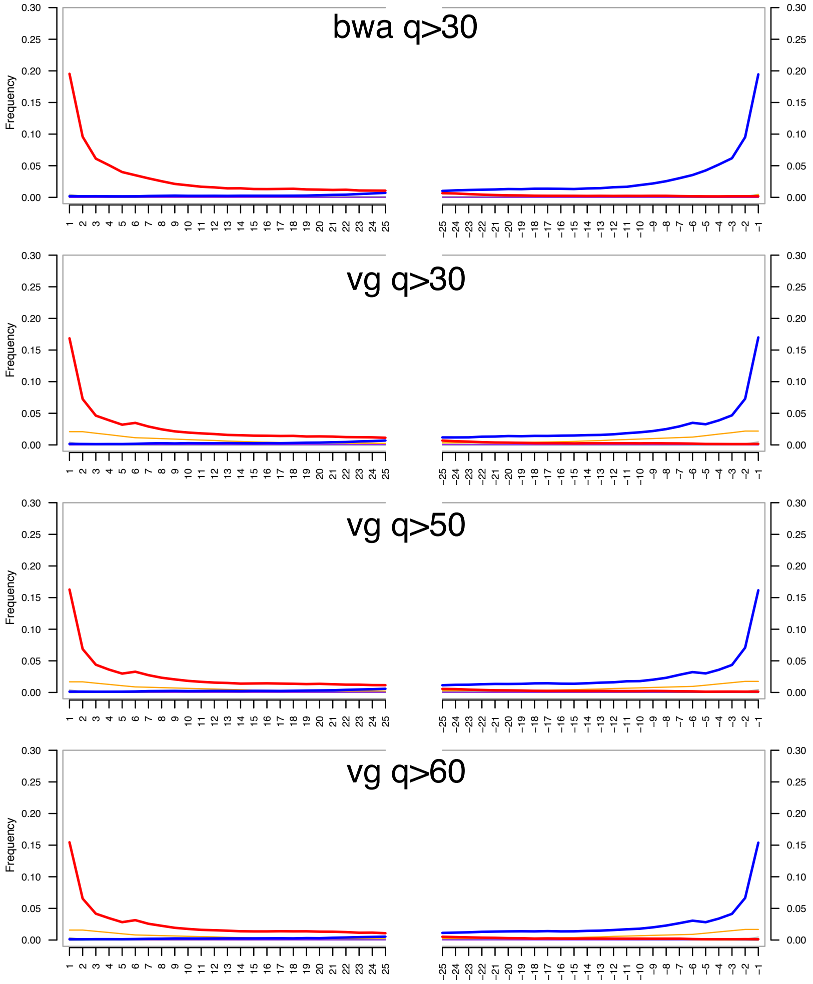

Figure S12: Comparison between deamination patterns in sequence read data belonging to a high-coverage Yamnaya sample processed with vg and bwa aln (-n 0.02) and filtered with different mapping quality thresholds. Red: C to T substitutions; blue: G to A substitutions, and orange: soft-clipped bases.

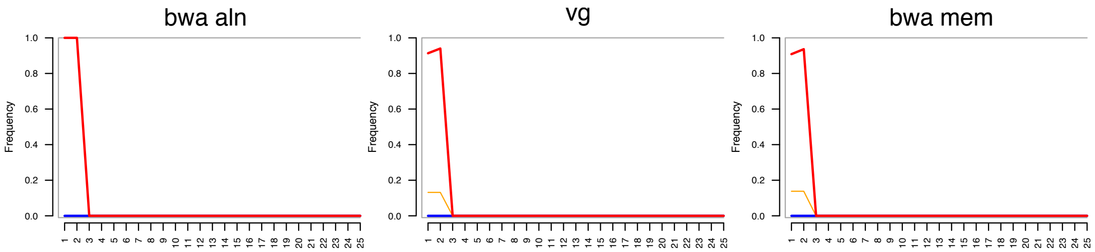

Figure S13: Comparison of deamination and softclipping patterns in sequences with artificially introduced C to T changes in the 1st and 2nd nucleotides of the 5'-end of sequencing reads. Softclipping (orange curve) is more frequent in vg and bwa mem, leading to a slight reduction in the red curve which indicates C to T changes.

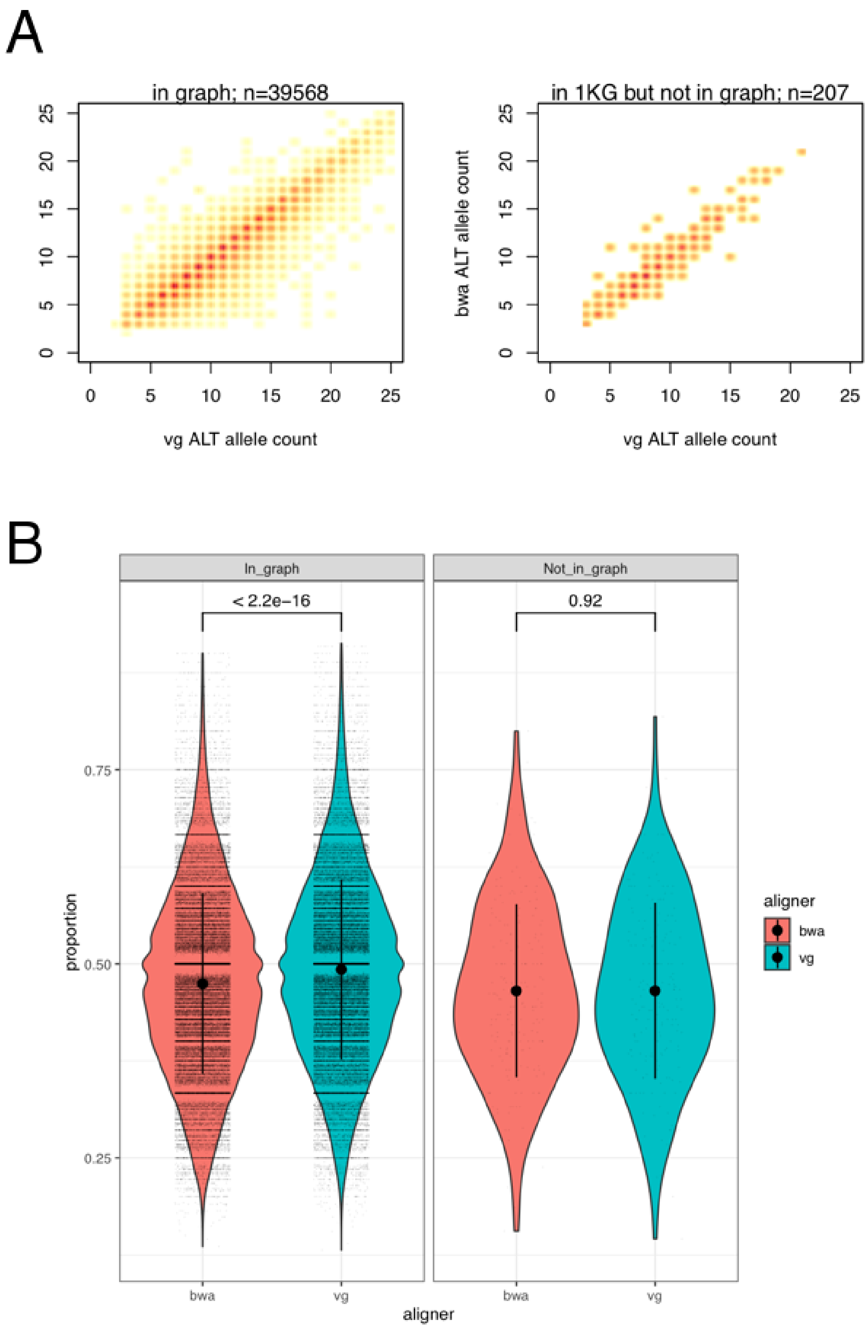

Figure S14: Comparison of alternate allele support in heterozygous transversions of chr1 called from the Yamnaya sample aligned with `vg map` and `bwa aln -n 0.02` . A) Alternate allele counts in `bwa aln` and `vg map` aligned samples at SNPs present (left) or absent (right) in the graph. B) Alternate allele proportion in `vg map` and `bwa aln` aligned bams at SNP sites present (left) or absent (right) in the graph.

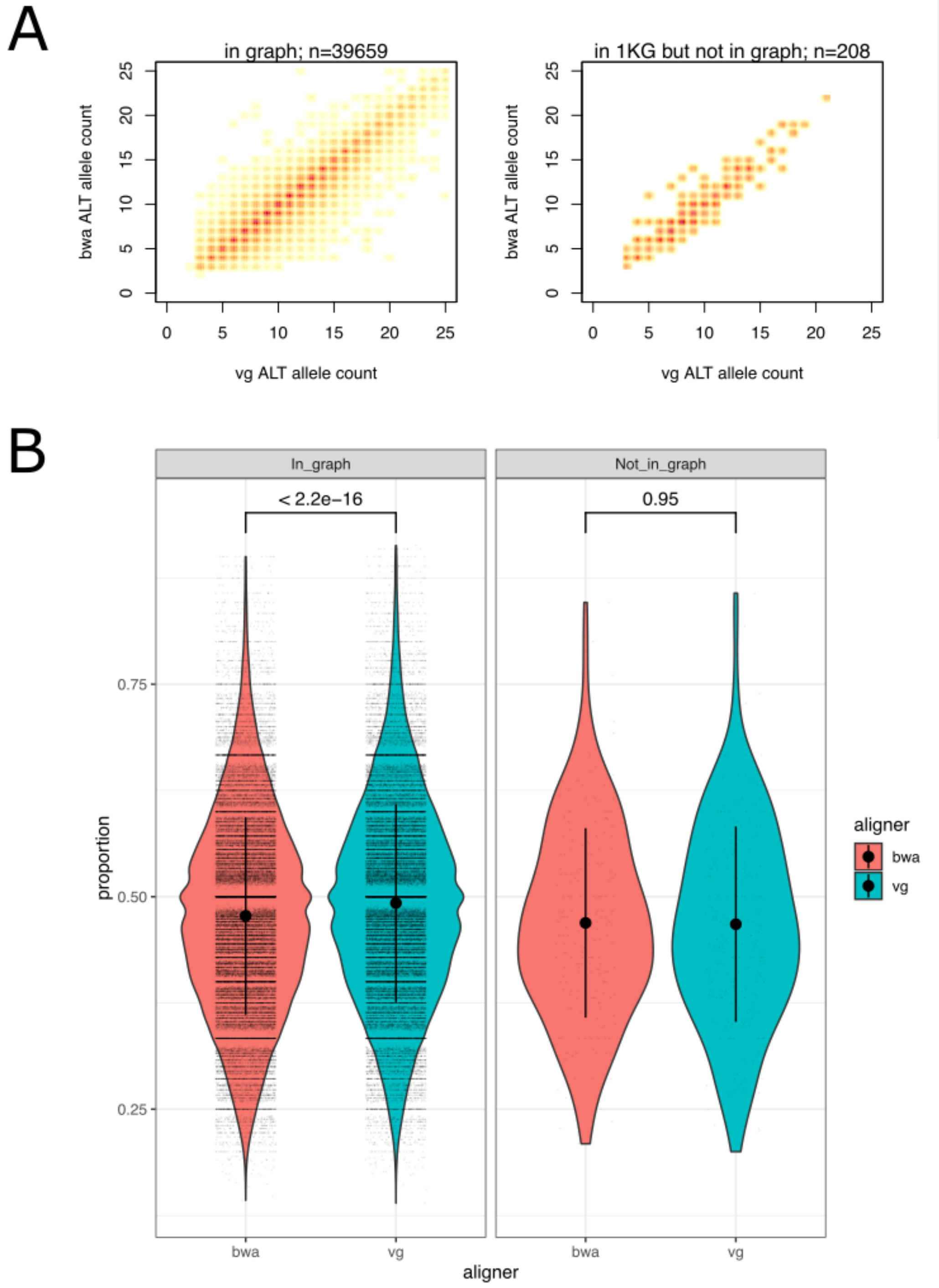

Figure S15: Comparison of alternate allele support in heterozygous transversions of chr1 called from the Yamnaya sample aligned with `vg map` and `bwa aln -n 0.01 -o2` . A) Alternate allele counts in `bwa aln` and `vg map` aligned samples at SNPs present (left) or absent (right) in the graph. B) Alternate allele proportion in `vg map` and `bwa aln` aligned bams at SNP sites present (left) or absent (right) in the graph.

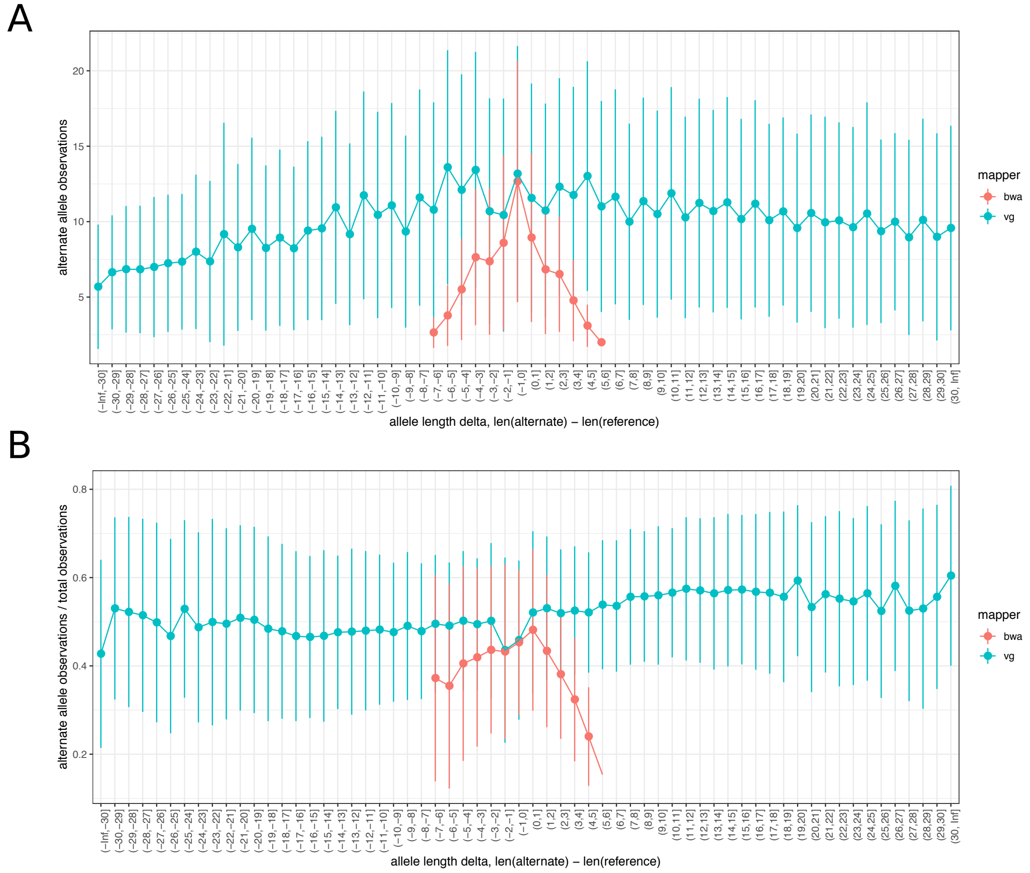

Figure S16: Comparison of indel calling between the Yamnaya sample processed with vg graph and bwa aln (-n 0.02) to demonstrate the relationship between allele length and reference bias. While with vg it is possible to call alternate alleles at indels of various lengths, bwa shows a strong reference bias to insertions and deletions greater than a few bp. A) Alternate allele observations at indels. B) Fraction of alternate alleles in all indel observations. Alleles were binned by their relative length to the reference allele.

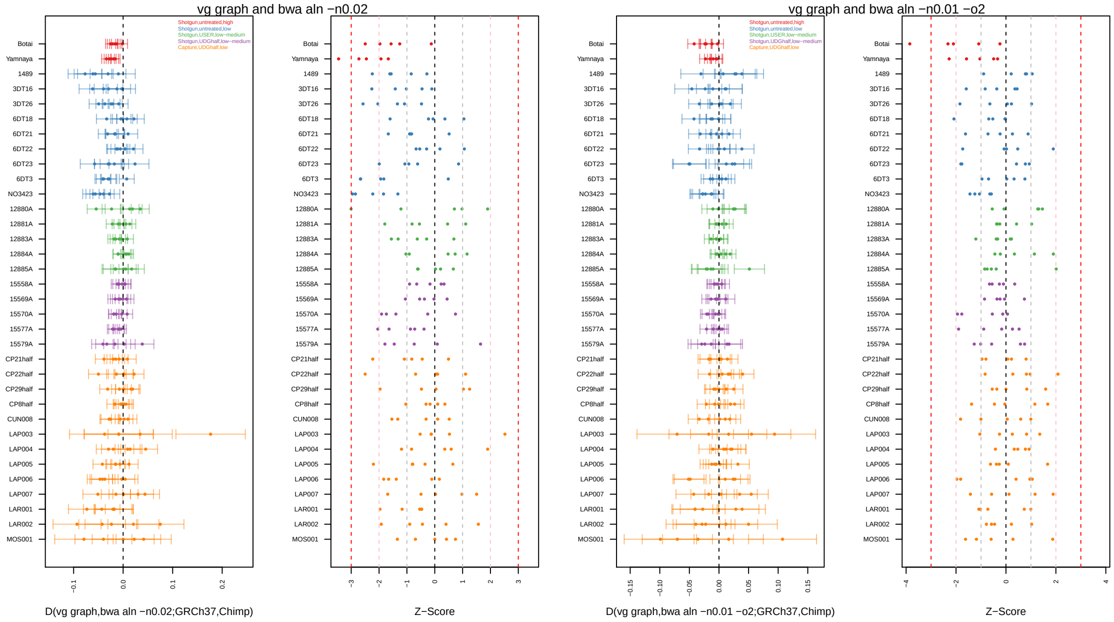

Figure S17: $D$-statistic test of reference bias comparing vg and bwa, using Chimp as an outgroup and including transversion SNPs only. The left two panels compare vg graph (mapQ50) and bwa aln -n0.02 (mapQ30) in terms of bias towards the reference and the right two panels compare vg graph (mapQ50) and bwa aln -n0.01 -o2 (mapQ25).

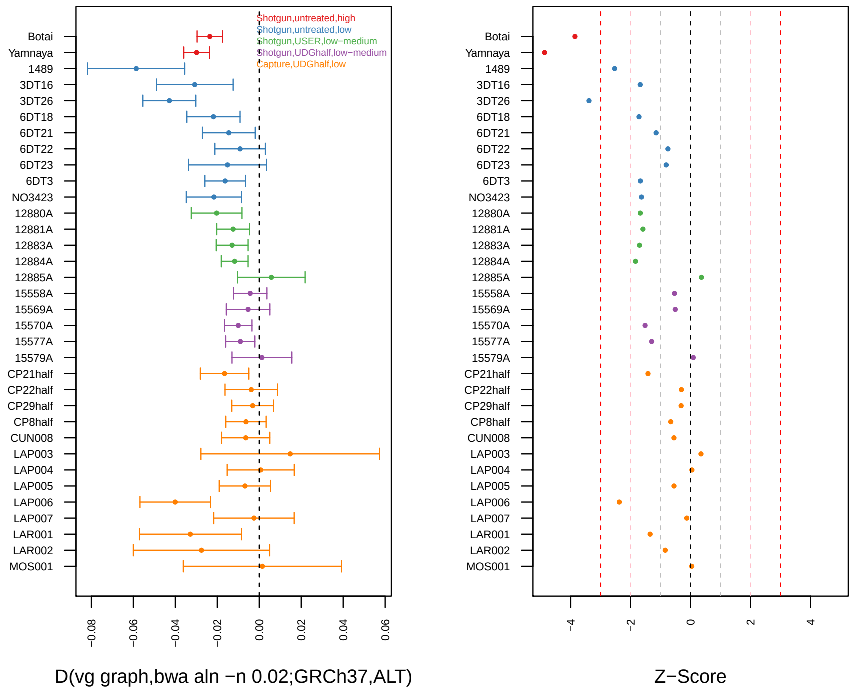

Figure S18: $D$-statistics of the form $D$(vg graph, bwa aln -n 0.02; GRCh37,alternate allele) estimated for 34 ancient individuals, using transversion SNPs only. The figure label indicates the type of sequencing, enzymatic treatment and genomic coverage for each sample.

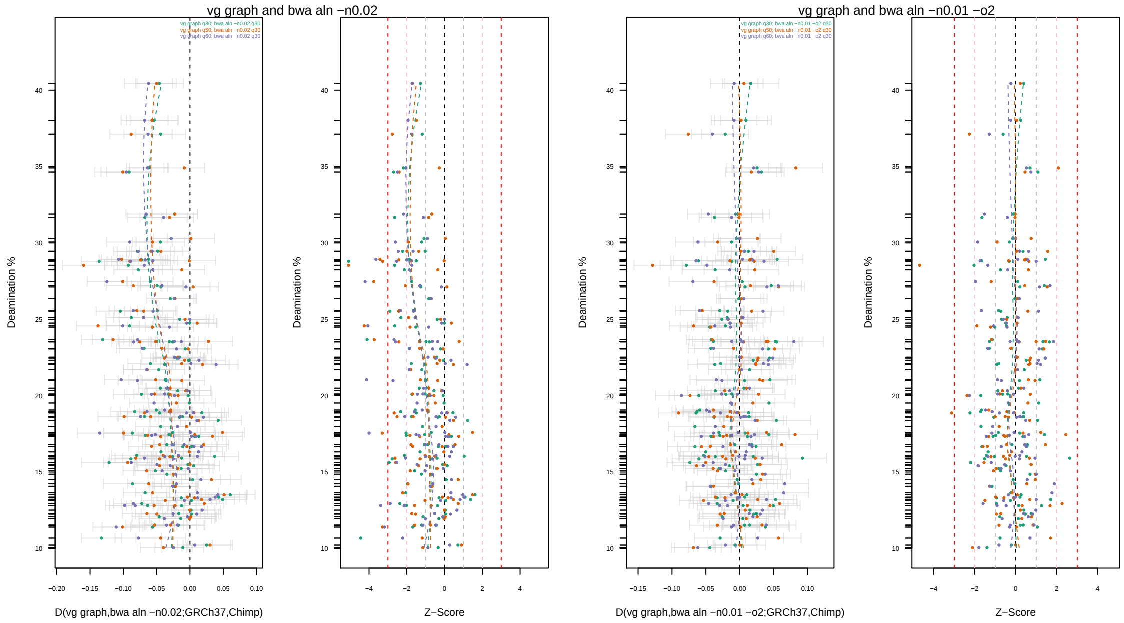

Figure S19: *D*-statistic based ABBA-BABA test of reference bias based on simulated data aligned with vg graph and bwa aln, and using Chimp as an outgroup and including transversion SNPs only. The left two panels compare vg graph (mapQ50) and bwa aln -n0.02 (mapQ30) in terms of bias towards the reference and the right two panels compare vg graph (mapQ50) and bwa aln -n0.01 -o2 (mapQ25).

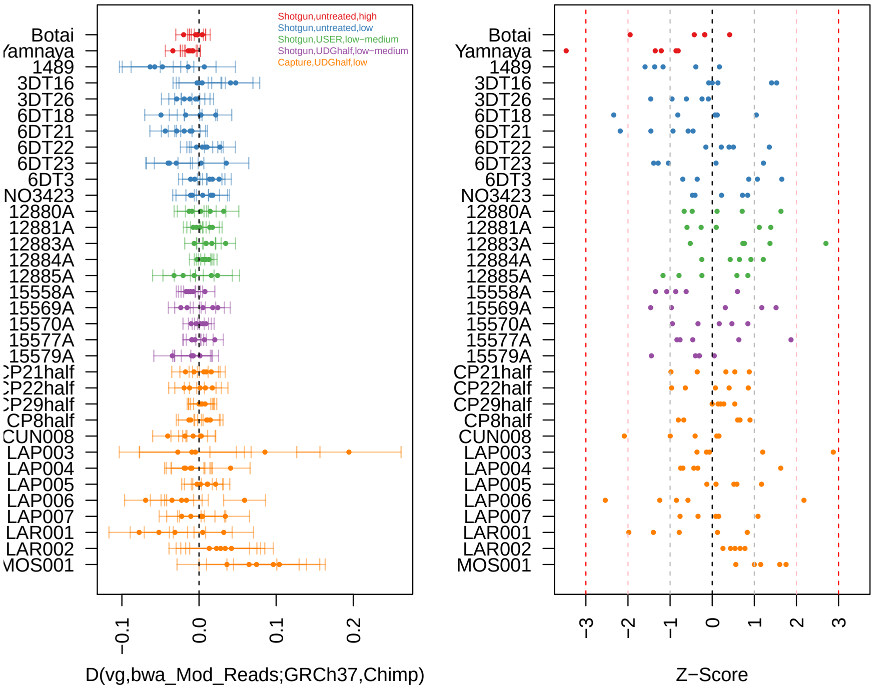

Figure S20: $D$-statistic based ABBA-BABA test of reference bias comparing bwa-modread and vg using Chimp as an outgroup and estimated with transversion SNPs only. We included 5 replicates per ancient sample to account for the randomness in pseudo-haploid genotype generation.

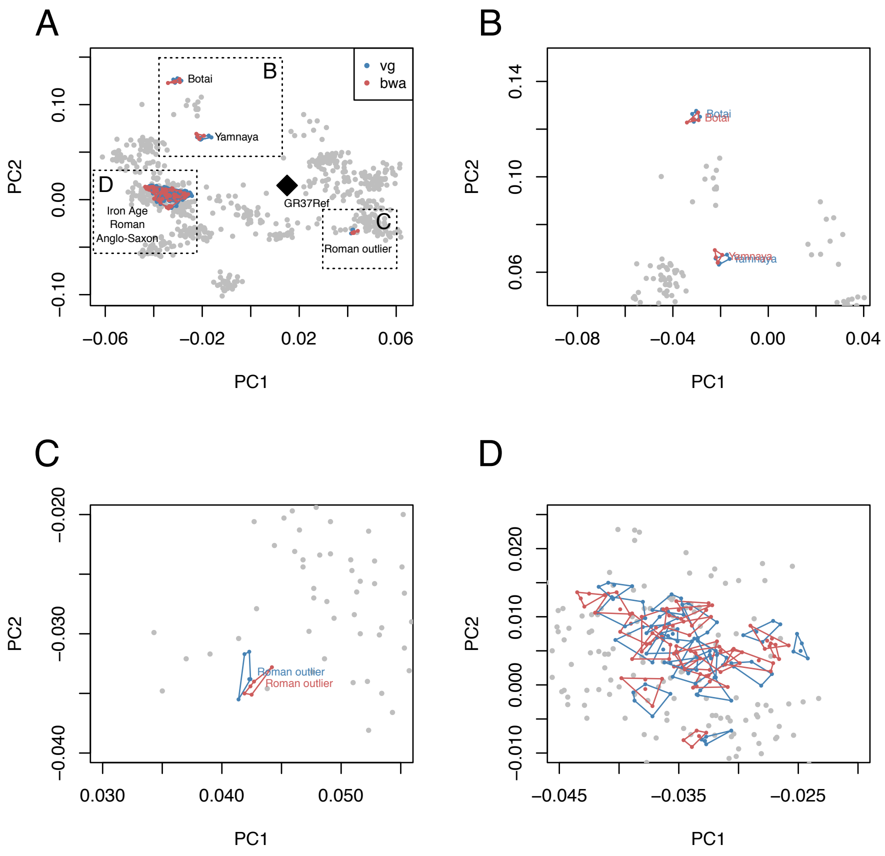

Figure S21: Principal Component Analysis estimated with present-day West Eurasians from the Human Origins dataset with ancient samples aligned with either vg graph (blue) or bwa aln (-n 0.02) (red) projected onto it. 5 replicates of random allele sampling for each one of the ancient samples were included in the analysis. A) Full view of the PCA. Zoomed in PCA regions: B) Botai and Yamnaya, C) Roman outlier, D) Iron Age, Roman and Anglo-Saxon samples.

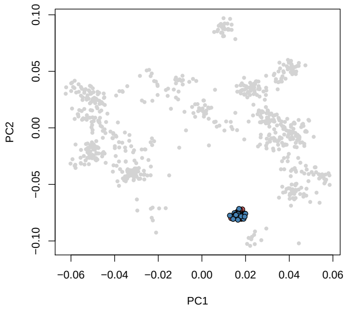

(a) Full PCA.

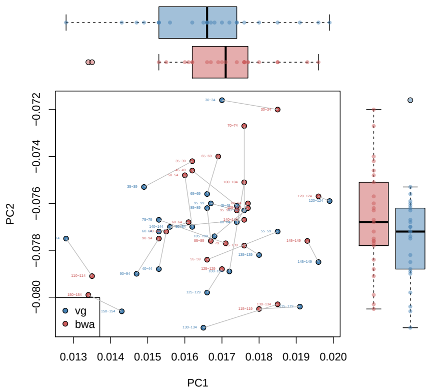

(b) PCA zoomed into the coordinates of the Yamnaya sample.

Figure S22: PCA estimated with West Eurasian populations from the Human Origins dataset and with the projected Yamnaya sample, aligned with either vg graph and bwa aln (-n 0.02), and divided into 25 different read length bins (30-155bp). Present-day populations are coloured in grey. Yamnaya pseudo-haploid genotypes obtained from vg (blue) or bwa aln (red) aligned reads. Each 5 bp bin is represented by a point. Genotypes called from the same read length bin in bwa aln and vg are connected by a grey line.

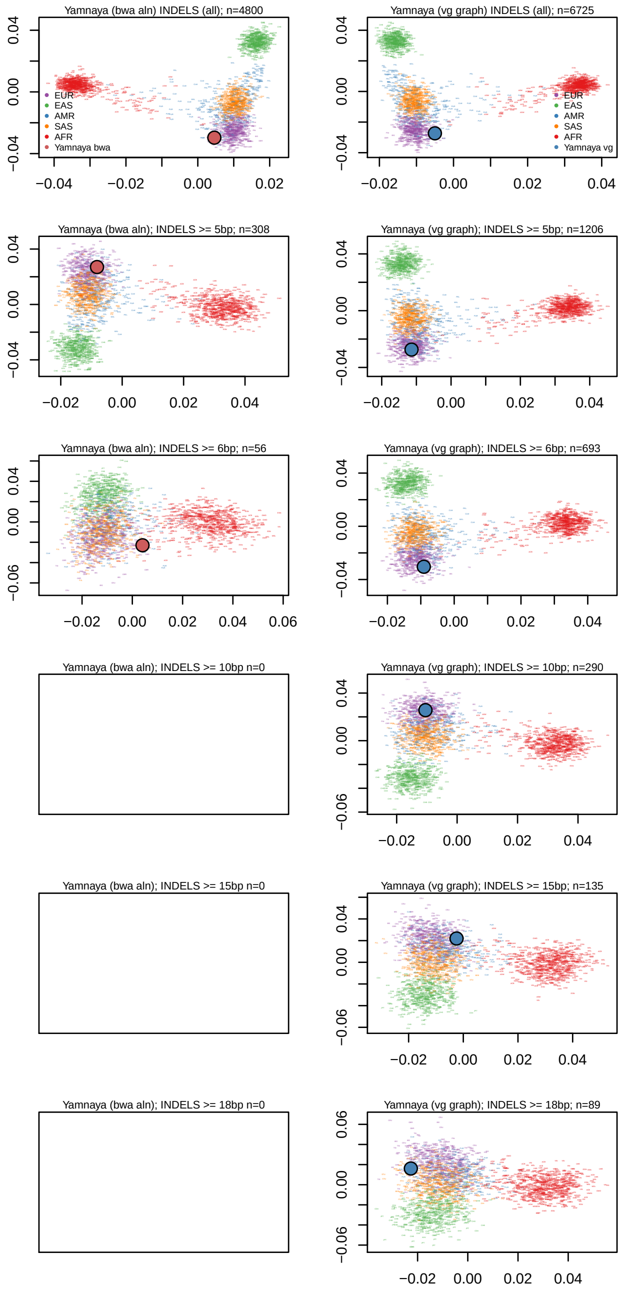

Figure S23: Principal Component Analysis estimated with chr21 indels from the 1000 Genomes project. The indels used for analysis were restricted to those identified either in Yamnaya aligned with bwa aln (-n 0.02,left) or with vg graph. While PCAs estimated with vg indels maintain a clear clustering of the 1000 Genomes superpopulations across a wide range of indel allele lengths, with bwa this is not the case, given it does not recover longer indels than a few basepairs. The x and y axes are PC1 and PC2. Some plots were intentionally left blank to demonstrate the inability of bwa to call indels greater than 6 basepairs.
